# Cis-regulatory elements within TEs can influence expression of nearby maize genes

**DOI:** 10.1101/2020.05.20.107169

**Authors:** Jaclyn M Noshay, Alexandre P Marand, Sarah N Anderson, Peng Zhou, Maria Katherine Mejia Guerra, Zefu Lu, Christine O’Connor, Peter A Crisp, Candice N. Hirsch, Robert J Schmitz, Nathan M Springer

## Abstract

Transposable elements (TEs) have the potential to create regulatory variation both through disruption of existing DNA regulatory elements and through creation of novel DNA regulatory elements. In a species with a large genome, such as maize, the many TEs interspersed with genes creates opportunities for significant allelic variation due to TE presence/absence polymorphisms among individuals. We used information on putative regulatory elements in combination with knowledge about TE polymorphisms in maize to identify TE insertions that interrupt existing accessible chromatin regions (ACRs) in B73 as well as examples of polymorphic TEs that contain ACRs among four inbred lines of maize including B73, Mo17, W22, and PH207. The TE insertions in three other assembled maize genomes (Mo17, W22 or PH207) that interrupt ACRs that are present in the B73 genome can trigger changes to the chromatin suggesting the potential for both genetic and epigenetic influences of these insertions. Nearly 20% of the ACRs located over 2kb from the nearest gene are located within an annotated TE. These are regions of unmethylated DNA that show evidence for functional importance similar to ACRs that are not present within TEs. Using a large panel of maize genotypes we tested if there is an association between the presence of TE insertions that interrupt, or carry, an ACR and the expression of nearby genes. TEs that carry ACRs exhibit an enrichment for being associated with higher expression of nearby genes, suggesting that these TEs may create novel regulatory elements. These analyses highlight the potential for TEs to rewire transcriptional responses in eukaryotic genomes.

**Data Availability:** In this study we utilize previously published datasets that are available through the following accessions: SRX4727413, SRR8738272, and SRR8740852.

## Introduction

Transposable elements (TEs) are highly repetitive DNA sequences found in most genomes. Variable genome size between related species has been partially attributed to the accumulation of TEs (Michael and Jackson 2013). The maize genome is replete with TEs, having >80% of the ~2500Mb of genomic space being composed of repetitive sequence and 64% annotated as complete TEs (Schnable et al. 2009; Jiao et al. 2016). TEs can be classified into two main orders based on their transposition intermediate, Class I RNA retrotransposons which commonly proliferate through “copy and paste” transposition and Class II DNA transposons that generally move through a “cut and paste” mechanism (Wicker et al. 2007). Barbara McClintock referred to these repetitive sequences as “controlling elements”, encompassing their potential to impact and regulate genes (McClintock 1951). Transposition enables these TEs to move throughout the genome potentially influencing functional regions. TEs may insert into coding regions and cause direct influence on gene function, but also may insert into existing regulatory regions or create new regulatory elements resulting in altered gene expression.

One mechanism of TE influence on gene expression is through disruption of regulatory sequences. TEs in the maize genome are dispersed throughout the chromosome including gene-rich regions of chromosome arms (Schnable et al. 2009; Baucom et al. 2009). Due to this interspersion of genes and TEs, many TEs have the potential to influence expression of genes. DNA transposons have been shown to display preferential insertion into genic regions while retrotransposons appear to be more present in heterochromatic, gene poor regions of the genome (Bennetzen 2000). A DNA transposon, mPing in Oryza sativa was found to preferentially insert into the 5’ regions of genes (Naito et al. 2009). S-elements in Drosophila melanogaster have been found to insert into the 5’ regions of several members of the Hsp70 heat shock gene family (Maside et al. 2002). Another type of TE, known as miniature inverted repeat transposable elements (MITEs), often insert into the last exon of genes, which may cause more impact than ordinary intron insertions (Guo et al., 2017). MITEs have also been found to play an evolutionary role in altering gene expression through contributing regulatory elements (Wessler et al., 1995).

TEs not only have the potential to disrupt regulatory sequence, but can also introduce novel regulatory elements into new genomic locations (Feschotte 2008; Chuong et al. 2017). TE insertions can also result in changes in the location of regulatory elements relative to nearby genes (Zhao et al. 2018; Lu et al. 2019). It has been shown that TEs can impact gene expression through several examples in maize including teosinte branched 1 (tb1), a gene responsible for the branching in the maize progenitor, teosinte (Studer et al. 2011). The regulatory region of tb1 is within the intergenic space ~60kb upstream of the gene (Doebley et al. 1997; Clark et al. 2006; Briggs et al. 2007). An essential insertion of a retrotransposon Hopscotch acts as an enhancer of gene expression resulting in the branching differences between maize and teosinte (Studer et al. 2011). Similar examples are observed in other species as well. Jordan et al. (Jordan et al. 2003) reported that nearly a quarter of all promoters characterized in humans contain TE sequences. Another study focusing on human T-cells identified DNase hypersensitive sites significantly overlapped with annotated TEs suggesting the presence of cis-regulatory regions (Sheffield et al. 2013). The existence of regulatory regions within TEs could represent examples of regulatory elements that have evolved to solely regulate expression of the TE itself as well as examples in which the regulatory elements within the TE have been co-opted to regulate nearby genes (Chuong et al. 2017; Zhao et al. 2018).

The question of how TEs impact the genome has been considered from different perspectives since McClintock first discovered their existence. There are many examples in which detailed analyses of specific QTL have revealed the importance of TE insertions in creating altered gene expression (Zerjal et al. 2012; Zhang et al. 2012; Yang et al. 2013; Castelletti et al. 2014; Mao et al. 2015). There have been hints that certain families of TEs are associated with genes that exhibit stress-responsive expression (Makarevitch et al. 2015) and that many TEs exhibit dynamic, tissuespecific patterns of expression (Anderson et al. 2019b). There is evidence that a substantial number of accessible chromatin regions are found within TEs (Oka et al. 2017; Zhao et al. 2018; Lu et al. 2019) and in some cases these sequences can provide evidence for regulatory activity (Zhao et al. 2018).

In order to assess the mechanisms by which transposons might influence cis-regulatory elements it is important to have an understanding of putative regulatory elements and transposon variaiton among genotypes. The availability of genome-wide identification of accessible chromatin regions (ACRs) in B73 (Ricci et al. 2019) and high-quality information on shared and polymorphic TEs (Anderson et al. 2019a) provides new opportunities to address the potential impact of TEs on gene regulation in maize. We characterized hundreds of examples of B73 ACRs that are interrupted by a TE insertion in another genotype and thousands of examples of ACRs that are within annotated TEs. TE insertions into ACRs can result in chromatin changes to the ACR in addition to the genetic change. Many of these ACRs within TEs show evidence of functional enrichment. Through analyses of putative regulatory regions and TE polymorphisms we can begin to evaluate how TEs may contribute to natural variation for gene expression in maize.

## Methods

### Annotation of Genes and TEs

Whole genome assemblies for B73 (Zm00001d) (Jiao et al. 2016), W22 (Zm00004b) (Springer et al. 2018), Mo17 (Zm00014a) (Sun et al. 2018), and PH207 (Zm00008a) (Hirsch et al. 2016) were used for genome-wide analyses. All analyses were done on assemblies of chromosomes 1-10 while all scaffolds were disregarded due to the inability to assess these regions across genotypes. Filtered structural TE annotations (Stitzer et al.; Anderson et al. 2019a) were used.

### Polymorphic TEs

Shared and non-shared TEs across genotypes were defined previously (Anderson et al. 2019a). Briefly, identification of shared and non-shared elements was determined through pairwise comparison between four maize inbred lines (B73, W22, PH207, and Mo17). Search windows were defined by the closest, non-overlapping genes to the query TE with a syntelog in the genome being assessed. For comparison, 400bp flanking tags were extracted for each annotated TE in the genome (for each genome assessed) centered at the start and end coordinates. These flank tags were mapped to the other genomes with use of BWA-MEM (Li and Durbin 2009) in paired-end mode. Further characterization was performed on those elements with tags mapped completely within the search window. Non-shared site-defined TEs were defined by alignment of only the outer 200bp of the flank tags where the distance between tags was less than twice the TSD length for the superfamily. This resulted in a total of 69,292 non-shared site-defined elements across all pairwise comparisons used for analyses (Anderson et al. 2019a).

A total of 509,629 non-redundant TEs defined in at least one of B73, Mo17, PH207 or W22 structural TE annotations were assigned as present or absent in 509 of the WiDiv inbred genotypes (Hansey et al. 2011). Two points of reference, 10 bp over left and right inner edges of a TE, were used to determine TE status in a particular genotype. TEs with a coverage >= 8 across both inner edges were classified as present while TEs with coverage < 7 across both inner edges were classified as absent. All other TEs were classified as ambiguous.

### Methylation data

In this study we utilized previously generated WGBS data for B73 seedling shoot, PH207 seedling shoot, Mo17 seedling leaf and W22 seedling leaf. Trim_glore(Martin 2011) was used to trim adapter sequences and read quality was assessed with the default parameters and paired-end reads mode. Reads that passed quality control were aligned to the B73v4 genome (non-B73 genotypes were also aligned to their corresponding genome assemblies). Alignments were conducted using BSMAP-2.90(Xi and Li 2009), allowing up to 5 mismatches and a quality threshold of 20 (-v 5 – q 20). Duplicate reads were detected and removed using picard-tools-1.102 (“Picard Tools – By Broad Institute”) and SAMtools (Li et al. 2009). Conversion rate was determined using the reads mapped to the unmethylated chloroplast genome. The resulting alignment file, merged for all samples with the same tissue and genotype, was then used to determine methylation level for each cytosine using BSMAP tools. Methylation ratio for 100bp non-overlapping sliding windows across the B73v4 genome in all three sequence contexts (CG, CHG, and CHH) was calculated (#C/(#C+#T)). Each 100bp window was categorized as methylated (>=40%), intermediate (20-40%), or unmethylated (<=20%) based on the CHG methylation level.

### ATAC-seq data

In this study we utilized previously generated seedling shoot ATAC-seq data for B73 (Ricci et al. 2019). Raw reads were trimmed with Trimmomatic v0.33. Reads were trimmed for NexteraPE with a maximum of two seed mismatches, palindrome clip threshold of 30, and simple clip threshold of 10. Reads shorter than 30 bp were discarded. Trimmed reads were aligned to the Zea mays AGPv4 reference genome 44 using Bowtie v1.1.147 with the following parameters: “bowtie –X 1000 -m 1 -v 2 --best –strata”. Aligned reads were sorted using SAMtools v1.3.1 and clonal duplicates were removed using Picard version v2.16.0 (http://broadinstitute.github.io/picard/).

### Identification of accessible chromatin regions (ACRs)

MACS2 was used to define accessible chromatin regions (ACRs) with the “--keep-dup all” function and with ATAC-seq input samples (Tn5 transposition into naked gDNA) as a control. The ACRs identified by MACS2 were further filtered using the following steps: 1) peaks were split into 50 bp windows with 25 bp steps; 2) to quantify the accessibility of each window, the Tn5 integration frequency in each window was calculated and normalized with the average integration frequency across the whole genome to generate an enrichment fold value; 3) windows with enrichment fold values passing a cutoff (25-fold) were merged together by allowing 150 bp gaps; 4) to remove possible false positive regions, small regions with only one window were filtered for lengths > 50 bp. The sites within ACRs with the highest Tn5 integration frequencies were defined as ACR “summits”.For the functional analysis of SNP, HiChIP, STARR-seq and eQTL data we utilized the same methods as described in Ricci, Lu, Ji et al., 2019. The difference lies in the subset of data that was used to focus on TE ACRs versus non-TE ACRs opposed to all distal ACRs in the genome.

### Determination of TE-ACR overlap

TE-ACRs were defined by an overlap of B73 ACR coordinates with the structural TE annotation coordinates. Each ACR was assigned to a single TE using bedtools closest based on the disjoined TE coordinates file. For those with a partial overlap of multiple TEs the ACR was assigned to the TE with the greatest overlap. Complete overlaps were defined by >80% of the ACR length overlapping a TE.

### Identifying TE-insertions into ACRs

Site-defined TE polymorphisms with the TE present in Mo17, W22, and/or PH207 and absent in B73 were utilized to identify TE insertions into ACRs. Bedtools intersect was run with all defined B73 ACRs and the site-defined insertions, using the B73 insertion site coordinates. Any site-defined TE in Mo17, PH207, and/or W22 that had an insertion site within the coordinate range of a B73 ACR was characterized as a TE-insertion into an ACR for further analyses.

### Analysis of methylation at TE insertion sites

Methylation for each TE insertion was defined for the TE present genotype (Mo17, PH207, or W22) and the TE absent genotype (B73). Changes in methylation were identified by comparing 100bp bin CG methylation of the ACR in B73 to CG methylation levels flanking the insertion site in the genotype present for the TE. The position of the insertion was determined by its location in the ACR by quartiles with the 1st and 4th quartile being insertions at the edge of the ACR and the 2nd and 3rd quartiles defined as insertions into the middle of the ACR.

### Analysis of Methylation at ACRs across genotypes

Gene anchor files have been one to one gene syntelogs pairwise between B73, Mo17, PH207, and W22. Gene key files are available at https://github.com/SNAnderson/maizeTE_variation and were filtered to only one-to-one gene matches. Bedtools closest upstream and downstream, ignoring overlaps, was run for each B73 ACR relative to gene anchor files between B73 and PH207, W22, and Mo17. The search window was defined by the closest upstream and downstream syntelog pair. BLAST was run for each B73 ACR sequence to PH207, W22, and Mo17 to identify sequence similarity in the search window for the corresponding genotype. The sequence coordinates were identified and bedtools overlap was run against the 100bp WGBS data for that genotype. The methylation state of the B73 ACR was compared to the methylation levels of the matching sequence in PH207, W22, and Mo17 (based on WGBS data aligned to the corresponding genome assembly). The ACR was characterized as methylated if the average level of CHG methylation was greater than 40% and unmethylated if the average level of CHG methylation was less than 20%. A change in methylated was identified by an ACR characterized as unmethylated in B73 having a methylated state in another genotype.

### Gene expression analyses

RNAseq datasets Hirsch et al. (Hirsch et al. 2014) and Kremling et al. (Kremling et al. 2018) were used to assess expression levels across 284 genotypes and 8 tissues (Table 2). To assess gene expression variation, the closest gene to each TE was determined in B73 and the expression of that gene was associated with the presence or absence of the TE in each of the 284 genotypes. Each element containing an ACR or inserting into an ACR was assigned to the closest B73v4 annotated gene (in either direction) using bedtools closest. Only one assignment was given for each TE and any TE annotated as containing the full sequence of a gene was removed from the analysis. For those with distal ACRs, HiChIP data was used to assign the gene if an interaction was identified (Table S2/S3). TE presence impact was determined for each TE-gene pair by averaging the expression values for TE-present genotypes and TE-absent genotypes and the log2(present/absent) value was calculated. To account for biases in the number of genotypes with each TE as present or absent a t-test was performed to determine the p-value for each gene in each tissue.

**Table 1:** B73 ACRs majority overlapping (>80%) or partially overlapping (<80%) annotated TEs

|  | Genic | Proximal | Distal |
| --- | --- | --- | --- |
| Total | 12587 | 9,183 | 10,651 |
| LTR | 138 (93) | 130 (94) | 1428 (225) |
| TIR | 25 (382) | 72 (387) | 63 (376) |
| Helitron | 301 (90) | 203 (74) | 433 (76) |
| Total TE | 464 (565) | 405 (555) | 1924 (677) |

\* values in ( ) represent partial overlaps (< 80%)

**Table 2:** RNA-seq and TE PAV dataset summaries

| Dataset | # Tissues | Genotypes w/ TE calls | TE-Insertion (N=377) |  | TE-ACR (N=2182) |  |
| --- | --- | --- | --- | --- | --- | --- |
|  |  |  | Significant (+/-) | Outliers (+/-) | Significant (+/-) | Outliers (+/-) |
| **Kremling et al. (2018)** | GRoot | 91 | 2 / 1 | 16 / 17 | 51 / 2 | 214 / 59 |
|  | GShoot | 91 | 3 / 10 | 0 / 27 | 55 / 4 | 204 / 54 |
|  | Kern | 84 | 4 / 3 | 15 / 23 | 67 / 2 | 240 / 60 |
|  | L3Base | 87 | 2 / 4 | 19 / 25 | 54 / 4 | 197 / 65 |
|  | L3Tip | 86 | 5 / 1 | 19 / 22 | 44 / 6 | 281 / 60 |
|  | LMAD | 54 | 3 / 0 | 17 / 27 | 30 / 8 | 265 / 86 |
|  | LMAN | 94 | 0 / 3 | 14 / 32 | 52 / 11 | 256 / 73 |
| **Hirsch et al. (2014)** | Seedling | 230 | 1 / 2 | 2 / 7 | 57 / 14 | 105 / 22 |
| **Non-redundant sum** | All of the Above | 259 | 9 / 12 | 57 / 86 | 153 / 37 | 667 / 295 |

## Results

To assess potential impacts of TEs on putative regulatory regions in the maize genome, we used a set of 32,421 previously characterized maize ACRs identified using an Assay for Transposase-Accessible Chromatin with sequencing, hereafter known as ATAC-seq (Ricci et al. 2019). Roughly similar numbers of ACRs were found within genes (12,587), proximal regions (within 2kb of genes – 9,183), and distal regions (>2kb form nearest gene – 10,651). Ricci, Lu, Ji et al (2019) documented evidence to support the functional relevance of distal ACRs through enrichment of genetic variants underlying morphological and expression variation (eQTL and GWAS), chromatin-chromatin (HiChIP) interactions, and self-transcribing active regulatory region sequencing (STARR-seq) enhancer activity. We sought to investigate the role that TEs might play in regulating gene expression by disrupting ACRs within the maize genome or in carrying ACRs within TEs (Figure 1A/B). To monitor TE insertions within TE-ACRs, we focused on the set of ACRs identified within the B73 genome (Ricci et al. 2019) and documented the TE insertions in these regions within the W22, Mo17 or PH207 genomes (Figure 1C). The TEs that contain an ACR (>80% of ACR within the TE) were determined by comparing the coordinates of ACRs within the B73 genome with the B73 TE annotations (Figure 1D). The set of TE insertions into ACRs and TEs containing ACRs were further characterized to understand how these changes might influence chromatin states and regulation of nearby genes.

**Figure 1:**
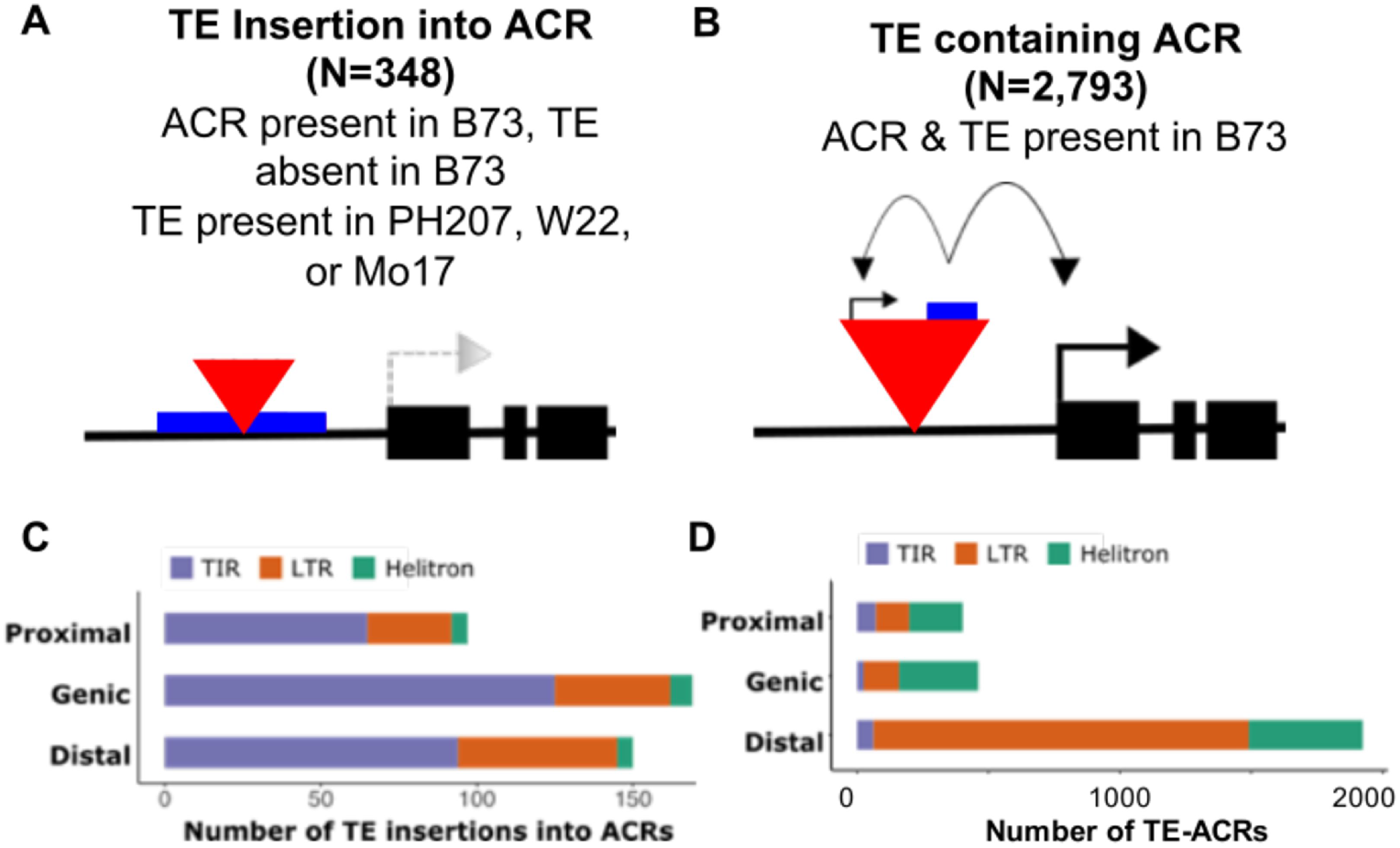
An overlap of TEs and accessible chromatin regions (ACRs). Schematic representation of the identified ACRs (blue) in the B73 maize inbred line and their interaction with TEs (red) and the potential impact on nearby genes. A) B73 ACRs that have a site-defined TE insertion in Ph207, Mo17 or W22. B) B73 ACRs that are found within B73 TE sequence. C) The number of TE insertions (as shown in A) in PH207, Mo17, or W22 into each ACR category (characterized by their position relative to annotated genes as genic, proximal, or distal) of ACR based on site-defined insertion sites in B73. Colors represent TE order. D) Number of TE-ACRs (as shown in B) by location relative to genes and TE order.

### Identification of TE insertions into ACRs

Of the 348 non-redundant instances of TE insertions into B73-defined ACRs, 176 TE insertions were found in Mo17, 82 insertions in PH207 and 158 insertions in W22. To determine the number of TE insertions expected by chance, we used a random set of control regions with similar size distribution as the ACRs. We observe significantly (Fisher’s exact p-value – 4.286e-07) more TE insertions in ACRs compared to the control regions (Figure S1A). The TEs that inserted were primarily terminal inverted repeat (TIR) DNA transposons with fewer examples of long terminal repeat (LTR) retroelements and Helitrons (Figure 1, Figure S1B). Several TIR elements have been found to preferentially insert within accessible chromatin (Jiang and Wessler 2001; Han et al. 2013; Noshay et al. 2019). The insertions into ACRs are highly enriched for members of the DTA and DTM superfamilies (Table S1) of TIR elements (Figure S1C). The TE insertions located within ACRs tended to represent relatively young TEs based on LTR similarity (Figure S1D).

### TE insertions into ACRs can result in altered chromatin

The ACRs represent regions of accessible chromatin and also lack DNA methylation (Ricci et al. 2019). The insertion of a TE in another haplotype could result in not only a genetic change to the DNA sequence, but also to changes in chromatin modifications or accessibility. DNA methylation data was generated for the same tissue type used for ATAC-seq in both B73 and PH207. There are 82 examples of PH207 TE insertions within B73 ACR regions and these were used to investigate the frequency of DNA methylation presence within the region classified as an ACR in B73. Specifically, we assessed the frequency of DNA methylation gains on one (uni-directional), or both (bi-directional) sides of the TE insertion (Figure 2A). In many cases the insertion of a TE within an ACR does not result in increased methylation of the regions with homology to the B73 ACR (Figure 2B). However for 37% of the TE insertions within ACRs, there are DNA methylation gains in the haplotype with the TE insertion (Figure 2C). TE insertions that are located within the outer quartiles of the ACR often result in methylation gains only on one side of the TE and is often the region closer to the edge of the ACR (Figure 2D). These analyses were solely focused on TE insertions within the B73 defined boundaries of the ACR. An analysis of 257 additional TE insertions (present in PH207, Mo17, or W22) located within 200bp of the ACR (present in B73) identified 30 additional examples in which a TE insertion near an ACR was associated with DNA methylation gains within the ACR. Together these analyses suggest that a subset of the TE insertions within, or near, ACRs can result in changes to the DNA methylation state of the region and are likely associated with changes in chromatin accessibility.

**Figure 2:**
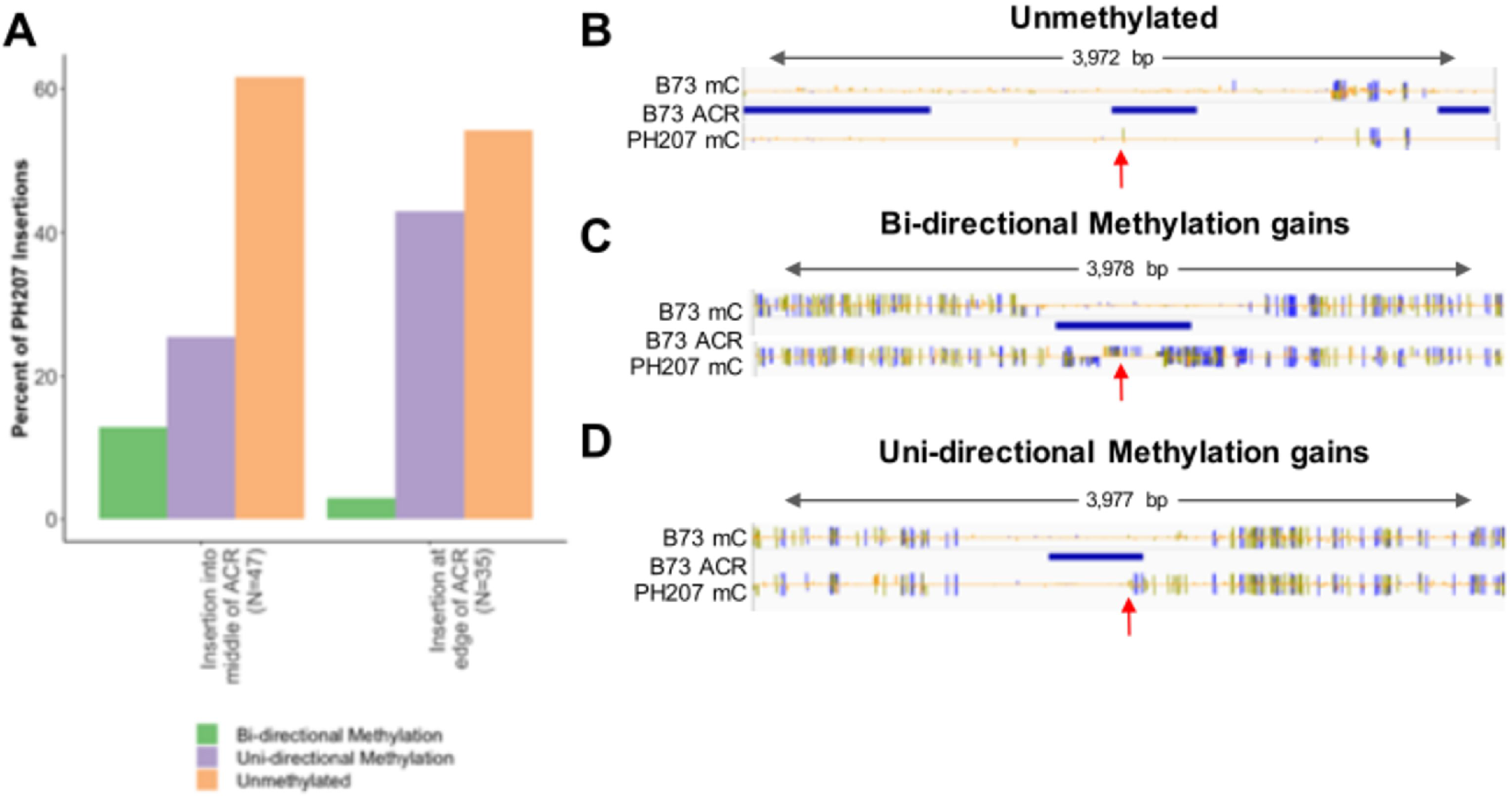
Methylation changes due to TE insertions in PH207. A) For every PH207 site-defined TE insertion into a B73 ACR the PH207 methylation status is defined as unmethylated (region remains unmethylated just as it was in B73), uni-directional methylation (methylation gain on one side of the insertion site), or bi-directional methylation (methylation gain on both sides of the insertion site). Insertions are broken into those that insert into the middle of an ACR (quartile 2 or 3) or those that insert into the edge of an ACR (quartile 1 or 4). WGBS data for B73 and PH207 were aligned to the B73 genome to visualize. IGV views display methylation level tracks (blue is CG, green is CHG, yellow is CHH), ACR region tracks, and TE insertion sites indicated by red arrows. These are shown for each methylation status; B) unmethylated, C) bi-directional methylation, D) uni-directional methylation.

### Identification of ACRs within TEs

In addition to the potential for TEs to disrupt existing ACRs, they also have the potential to carry sequences that lead to an accessible chromatin state and potentially move these sequences to new genomic locations (Figure 1B). We focused on characterizing examples of the ACRs that are identified in the B73 genome located within or overlapping annotated TEs. Of the 32,421 identified ACRs in maize, 4,590 have at least a partial overlap with an annotated TE (Table 1). It is worth noting that this is likely an underestimate of the number of true ACRs within TEs as the identification of ACRs relied upon uniquely mapping reads (Ricci et al. 2019). Many TEs are repetitive and have enough similarity to other family members to preclude uniquely mapping reads, which means that the number detected using unique mapping represents only a subset of actual accessible regions within TEs (Figure S4). In both leaf and ear tissue there is no evidence for enrichment of unique mapping reads in ATAC-seq data suggesting the presence of accessible chromatin within repetitive regions (Figure S4A). On a per-TE family basis, in which we could determine the number of reads that map to a family (both multiple mapping and unique mapping reads), there is evidence for some families with substantially more multi-mapping reads (Figure S4B). However, the multi-mapping reads cannot be attributed to a single genomic location and therefore we focused on the ACRs classified based on unique mapping reads for the remainder of our analyses.

Among the 4,590 TE-ACRs, there are 2,793 examples in which the majority (>80%) of the ACR is located within the TE and another 1,797 that have partial overlap (<80%) (Table 1; Figure S3A). These 1,797 partial overlaps may represent instances in which the ACR within the TE includes some adjacent sequence or may represent instances in which the TE inserted into an existing ACR and the accessible region spreads to encompass a portion of the TE. ACRs within TEs are more common for distal ACRs than for the other types of ACRs, especially for ACRs with majority (>80%) overlap with a TE (Figure S3A). The partial overlaps of ACRs with TEs have a high frequency of TIR elements, while the majority (>80%) overlap TE-ACRs have much higher frequencies of LTR elements (Figure S3A). Given the potential for the partial overlaps to represent instances of TE insertion into or near ACRs, rather than carrying the ACR within the TE, we focused on the majority (>80%) overlaps for the analyses of ACRs within TEs.

The 2,793 examples of majority TE-ACR overlap mostly (69%) comprise examples of distal ACRs (Figure 1D). Even though only 0.98% of all maize TEs contain an ACR, 19% of the distal ACRs are located within a TE (Table 1). Given an expectation that TEs would not contain accessible chromatin, this represents a large number of unexpected ACRs within TEs. However, if we assume that ACRs are randomly located in genomic sequence then the fact that 19% of distal ACRs are found within TEs is actually substantially fewer than expected (72% of random distal regions with size distribution similar to ACRs overlap a TE) given the amount of sequence attributed to TEs in the maize genome. The distal ACRs were further classified based on the patterns of several chromatin modifications into four groups; K-acetyl enriched, H3K27me3 enriched, transcribed and unmodified (Figure S3B) (Ricci et al. 2019). The TEs containing ACRs are enriched (chisquare p-value < 2.2e-16) for the transcribed class which is characterized by H3K4me3 and H3K36me3 along with acetylation marks and low DNA methylation levels similar to patterns seen in the promoters of expressed genes. This suggests that at least a portion of the ACRs found within TEs may represent promoters for expressed transposable element products. Prior work monitored expression of TEs in a variety of B73 tissues (Anderson et al. 2019b). Of the TEs containing an ACR classified as transcribed, 48% show observable expression levels in at least one tissue (Figure S3C). The TEs containing ACRs in the other classes (chromatin marked and unmodified) have lower frequencies of expressed elements, but are still expressed more often than non-ACR TEs (Figure S3C).

### Evidence for potential functional regulatory elements within TEs

Ricci, Lu, Ji et al., 2019 used several approaches to provide evidence for functional impacts of distal ACRs. Focusing on the 10,651 distal (>2kb from nearest gene) ACRs, we sought to determine whether there were differences in the support of functional impact for ACRs within TEs (TE-ACR) compared to ACRs located outside of TEs (nonTE-ACR). The frequency of SNPs is reduced within ACRs and this effect becomes even more pronounced when focusing on the TE-ACRs (Figure 3A). The analysis of the frequency of GWAS-associated SNPs revealed enrichment within both TE-ACRs and nonTE-ACRs (Figure 3B). TE-ACRs also show an enrichment for eQTL, although the level of enrichment is not as strong as observed for nonTE-ACRs (Figure 3C). The difference in the level of eQTL enrichment for TE-ACRs and nonTE-ACRs could be due to the differences in composition among the four chromatin classes of ACRs. The transcribed ACRs generally have lower enrichment than observed for some of the other classes (Figure S5). For ACRs to influence expression they would likely need to interact with nearby gene promoters. HiChIP analysis of chromatin interactions reveal similar enrichment for ACR-genic interactions for both TE and nonTE ACRs (Figure 3D-E). STARR-seq can identify sequences that can provide functional enhancer activity. STARR-seq analysis of maize accessible chromatin fragment activities in maize leaf protoplasts showed similar levels of enrichment for enhancer activity for TE and nonTE ACR sequences (Figure 3F).

**Figure 3:**
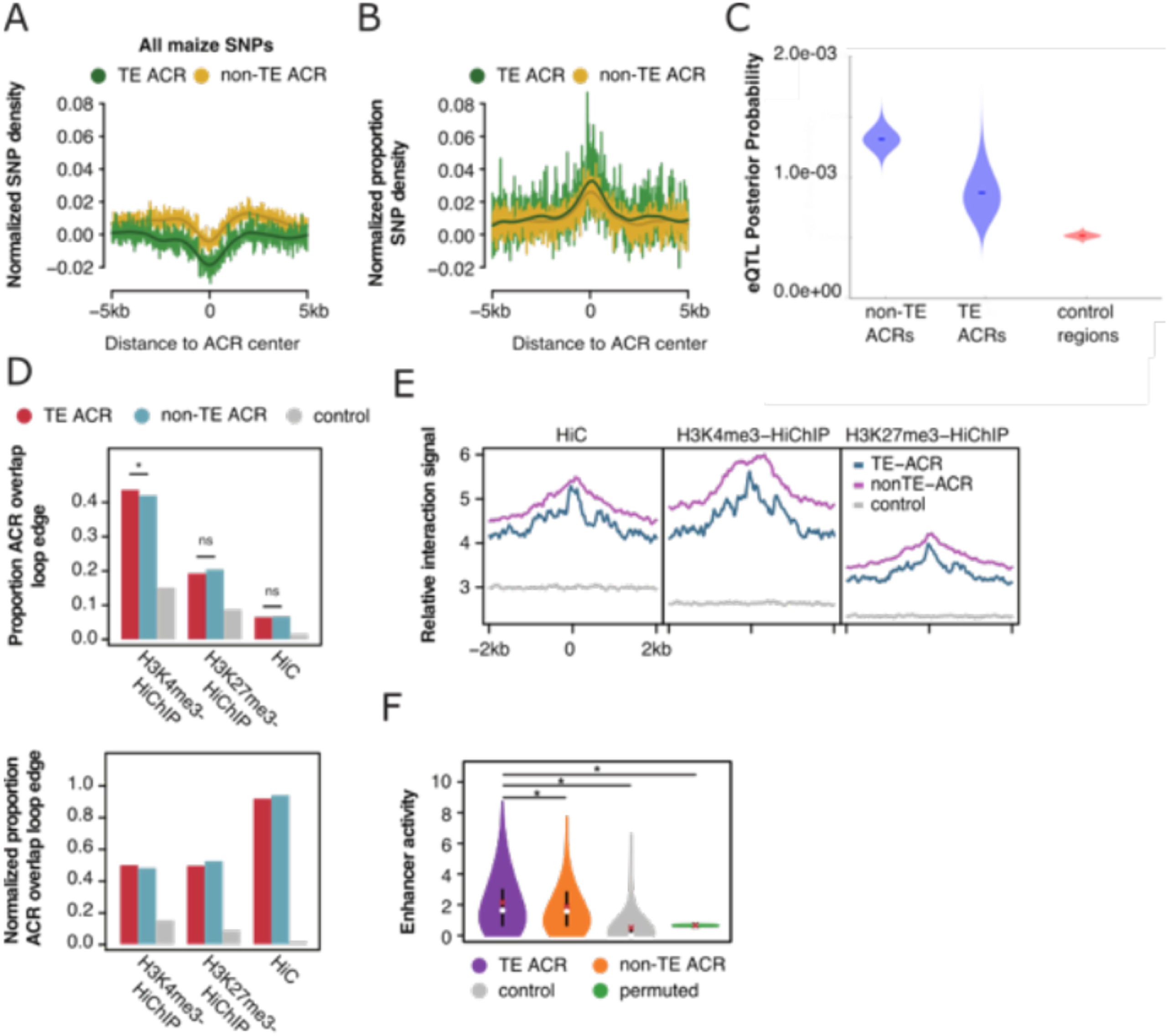
Functional differences between TE and non-TE accessible chromatin regions among distal ACRs. A) Normalized (control) SNP density among maize inbred lines averaged across 10kb regions centered on TE and non-TE ACRs. B) Proportion of GWAS hits (out of all maize SNPs) normalized by control enriched within 10kb windows centered on TE and non-TE dACRs. C) eQTL posterior probability for TE and non-TE ACRs compared to control regions. D) Contrasts between the proportions of dACRs overlapping an I-G loop between TE-ACRs and non-TE ACRs. Chi-square, *P-value < 0.05. E) Relative enrichment of chromatin interaction tags across 4kb windows centered on TE ACRs and non-TE ACRs across the three types of chromatin loops. F) Distribution of enhancer activities for dACRs split by the presence/absence of TEs, control regions (n=4,406) and the means of a permutation (10,000x). Statistical differences between TE and non-TE ACRs were evaluated with Mann-Whitney rank sum test. Statistical differences between distribution means and permuted regions were estimated as empirical P-values. ns, not significant; *P < 0.05

### Enrichment for certain TE families containing ACRs

TEs are classified into order, superfamily, and family based on transposition mechanism, structural components and sequence similarity. The ACRs that are located within TEs may represent TE family-specific properties in which multiple members of the same family contain an ACR or could represent instances in which the local chromatin neighborhood for a specific TE insertion allows the formation of an ACR. There are 356 of the 2,793 TE-ACRs that are located within single-member TE families. Among the remaining 2,437 TE-ACRs that are within multi-member TE families, 557 are only in one of the TEs in the family containing an ACR. This suggests that the majority of TE-ACRs are not a reproducible feature of the family members. A caveat to these results is the repetitive sequences which would not have been captured through the unique mapping ATAC-seq analysis and therefore additional members of a family may contain accessible chromatin regions (Figure S4B).

There are examples of TE-ACRs that are found in multiple members of a TE family. There are 112 TE families with at least two members with an ACR. There are only 10 of these families (with at least 3 elements) in which >30% of the elements have an ACR (Figure S6A). These examples of TE families with multiple members with ACRs were identified based on utilization of unique mapping reads. It is quite possible that additional members of these families may contain ACRs that were not identified because they are in regions that are highly similar in multiple TEs and therefore are multi-mapping. Two families in particular, RLX00813 and RLX01441, were found to display increased coverage when multi-mapping was allowed (Figure S6B).

### ACRs within TEs show variable DNA methylation patterns among genotypes

In general, TEs are considered to have quite high levels of DNA methylation, but ACRs typically lack DNA methylation (Oka et al. 2017; Lu et al. 2019; Ricci et al. 2019). The presence of ACRs within TEs led us to investigate the DNA methylation level of these sequences. We found that while TEs containing an ACR show quite high levels of DNA methylation throughout most of the TE, the ACR section is essentially unmethylated (Figure 4A-B). Visual inspection of several examples reveal that the ACR region represents a small window of unmethylated DNA within the largely methylated TE (Figure 4C-D).

**Figure 4:**
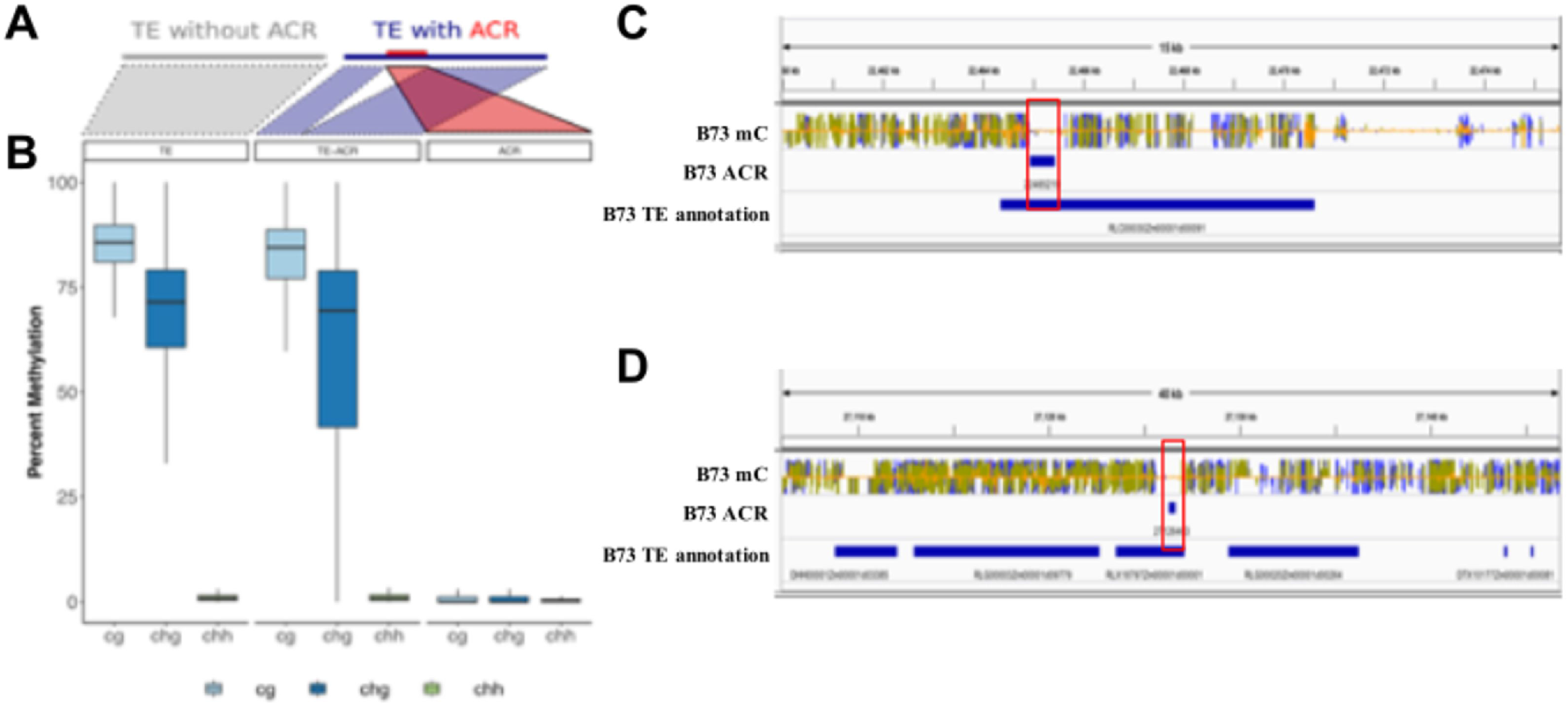
TE-ACR methylation patterns. A) Schematic representation of a TE without an ACR (grey) and a TE containing an ACR (blue) with the ACR sequence shown in red. B) Methylation levels of TEs without ACRs, TEs with an ACR (excluding ACR bins), and ACRs showing the trend that TEs maintain similar levels of high CG and CHG methylation with and without an ACR but the ~300bp region of an ACR is unmethylated. C/D) IGV view of TE with an ACR and the methylation levels (CG blue, CHG green, CHH yellow) over a majority of the TE and absence over the ACR

We hypothesized that the presence of an unmethylated region within a TE might be somewhat unstable and could be subject to changes in DNA methylation state among different haplotypes at a higher frequency than ACRs not located within TEs. An analysis was performed using a set of B73 ACRs that have a matching sequence at a syntenic location in PH207, Mo17, or W22 and have DNA methylation data available for both genotypes. These include ACRs within TEs that are present in both genomes and ACRs that are present in non-TE sequence (nonTE ACRs). While less than 3% of the nonTE ACRs exhibit gains of CG methylation across each of the genotypes, there are over 12% of the ACRs that are located within TEs that exhibit high levels of CG methylation (Figure 5A). Visual inspection of several loci suggest gains of both CG and CHG methylation over the full ACR sequence in these examples (Figure 5B-C). These observations suggest that ACRs within TEs may exhibit less stability among genotypes than ACRs in nonTE regions of genomes.

**Figure 5:**
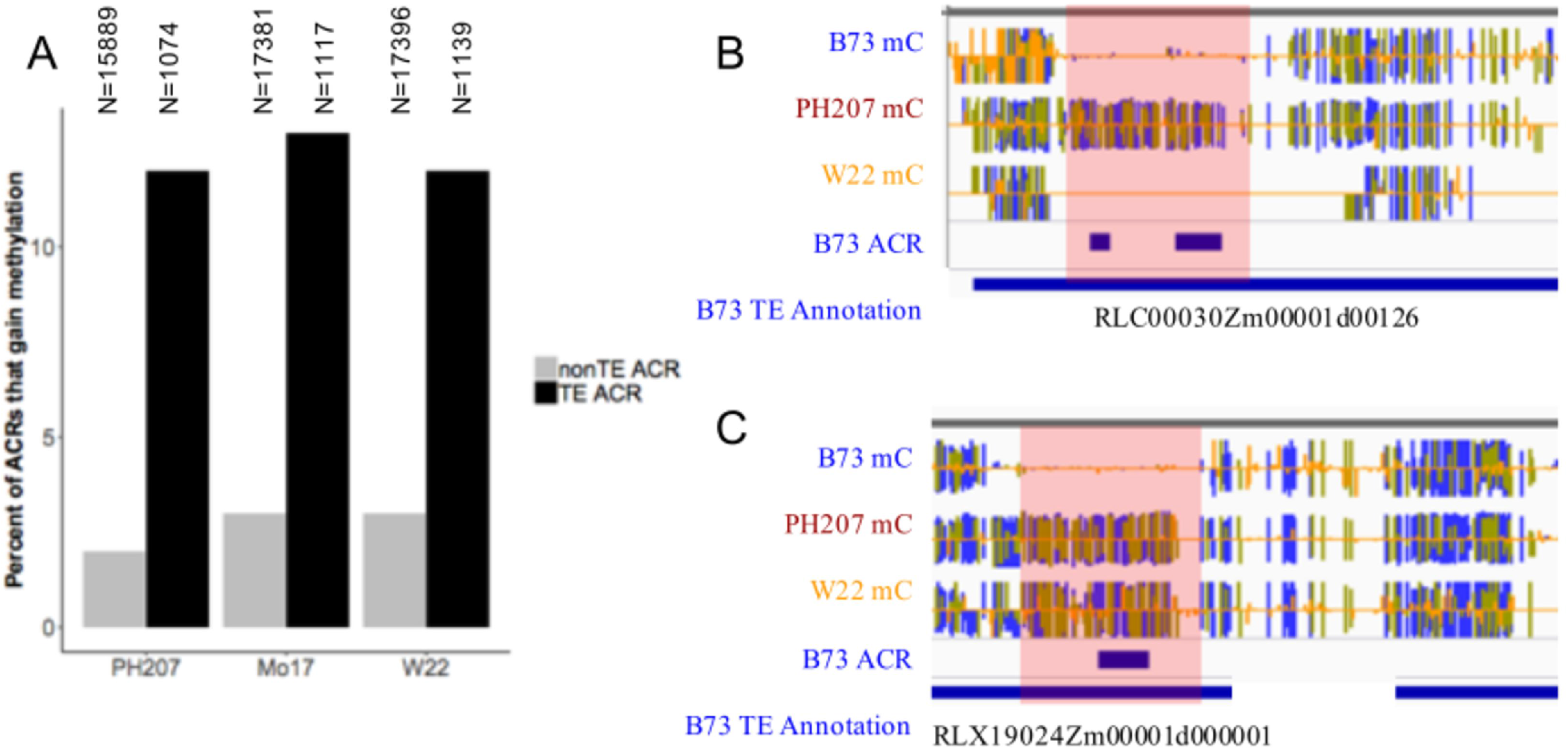
Unmethylated (open chromatin) regions in TEs are less stable than nonTE open chromatin regions. A) Percent of ACRs that gain methylation in PH207, Mo17, or W22 for non-TE ACRs (grey) and TE ACRs (black). B/C) IGV view of B73 TE annotation with unmethylated ACR in B73 and the same region as methylated in PH207 and/or W22. Methylation tracks show CG methylation in blue, CHG methylation in green, and CHH methylation in yellow.

### TE presence association with gene expression

Polymorphic TEs that interrupt an ACR or create novel ACRs in some haplotypes have the potential to influence the expression of nearby genes. To assess the potential for these polymorphic TE-ACR interactions to influence gene expression, we sought to associate the presence/absence of TEs with the changes in relative expression levels for nearby genes in panels of diverse germplasm. De novo assembled genome sequences of B73, Mo17, PH207 and W22 were used to generate de novo TE annotations in these four genomes (Anderson et al. 2019a). The presence or absence of these TEs was assessed in a larger (>500 inbreds) panel of diverse maize lines using alignments of whole-genome shotgun sequencing reads to the TE-flanking sequence junctions (see methods for details). This approach provides robust assignments of presence or absence for many genotypes but in some cases there is not clear evidence and the TE status is classified as ambiguous in that genotype. The TE polymorphism information was used to investigate variation in gene expression in several RNA-seq datasets (Hirsch et al. 2014; Kremling et al. 2018; Mazaheri et al. 2019). Each of these datasets included samples from a panel of genotypes that were collected at similar tissue stages.

Each polymorphic TE that disrupts a B73 ACR or provides an ACR in B73 was assigned based on HiChIP interactions or proximity to the nearest gene. TE-gene pairs where the gene is present completely within an annotated TE were disregarded for this analysis. We then assessed the difference in expression for genotypes with or without the TE insertion across the two datasets incorporating 284 genotypes and 8 tissues. (Table 2; Figure S8) allowing separate tests of potential associations between TE polymorphisms and expression level in multiple tissues. We initially focused on the set of 377 TE insertions into an ACR, which we hypothesized may result in reduced expression for the nearby gene. There are 21 instances (5.6% of all TE-gene pairs) in which we found a significant (q-value <0.05 and >2-fold-change) change in expression for the nearby gene (Table 2). These include 9 genes in which higher expression was observed for the haplotype containing the TE insertion, and 12 examples of lower expression when the TE is present. In 10 of the 21 significant associations, we found a significant association between the presence of the TE and expression levels in multiple tissues. In addition to the genes with significant associations, we also noticed that there is an apparent excess of many ‘outlier’ expression states for which the genotype with (or without the TE) has a >30-fold change in expression but there is limited statistical significance because one of the haplotypes is rare (Figure S8A). To determine if there is a significant excess of these outliers, we performed separate permutation tests in which the genotype-expression or genotype-TE presence classifications were randomized. These were separately performed for each of the expression datasets and were used to determine the number of significant or outlier expression changes expected by chance within this data structure (Figure 6A). The TE insertions into ACRs consistently exhibit more outliers than expected by chance with reduced expression of the haplotype with the TE present for each of the expression datasets (Figure 6A).

**Figure 6:**
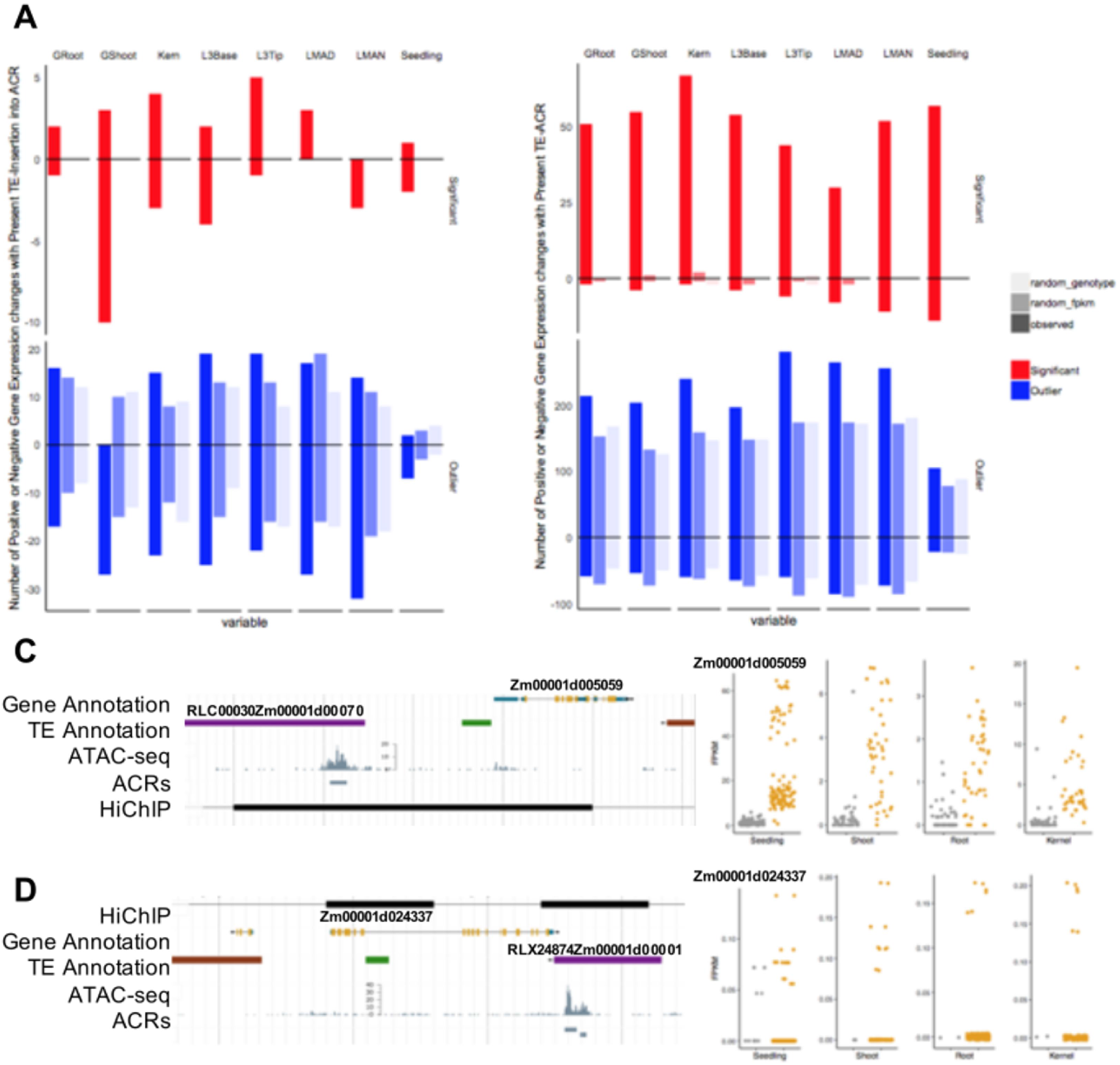
TE PAV association with gene expression. A) Number of TE-Insertions that result in significant (red) or outlier (blue) expression changes of nearby genes by tissue for observed and randomized genotype or randomized RNA-seq controls shown by shading. B) Number of TE-ACRs resulting in significant or outlier expression changes. C/D) Examples of significant gene expression changes associated with TE presence. Left: Genome browser view of the TE, Gene, and ACR. Right: Dotplot of gene expression for genotypes present (yellow) or absent (grey) for seedling, shoot, root, and kernel corresponding to the TE-Gene pair.

We next assessed the 2,182 polymorphic insertions of TEs containing ACRs near genes which were hypothesized to have positive influences on the expression of the nearby gene. There were 190 significant associations (8.7% of all tested TE-gene pairs) and 81% of these significant associations exhibit higher expression for the nearby gene (Figure S8B, Table 2). Many (49%) of the significant positive associations between the presence of the TE and the expression of the nearby gene were identified in multiple tissues while fewer (18%) of the negative associations were identified in multiple tissues. Figure 6C-D shows two examples of a TE located near a maize gene with significant positive associations with expression in multiple tissues. In both of these examples there are HiChIP interactions between the ACR within the TE and the nearby gene based on data from Ricci, Lu, Ji et al (2019). The permutations tests identify very few significant associations (Figure 6B). The analysis of rare outlier expression states also reveals an excess of positive associations in which the haplotype containing the TE exhibits a higher expression level (Figure 6B).

## Discussion

Many eukaryotic genomes show evidence for both recent amplification of transposable elements as well as turnover of elements through deletions (Bennetzen and Kellogg 1997). Insertions of transposons into genes or regulatory elements can lead to loss-of-function mutations which are presumed to be primarily deleterious. However, there is growing evidence that TEs may also contribute to re-wiring of transcription of nearby genes (Weil and Martienssen 2008; Feschotte 2008; Lisch 2013; Chuong et al. 2017). Transposon insertions that affect expression of a nearby gene are the molecular basis for allelic variation at several loci important for maize domestication and improvement (Studer et al. 2011; Yang et al. 2013; Castelletti et al. 2014). There are also examples in maize and other species in which transposon insertions may influence regulatory influences on nearby genes (Jiang et al. 2004; Cavrak et al. 2014; Makarevitch et al. 2015; Zhao et al. 2018). While specific examples have been identified, the genome-wide frequency for these TE influences has not been characterized. Advances in our knowledge of genome-wide TE polymorphisms (Stitzer et al.; Anderson et al. 2019a) as well as the identification of proximal and distal putative cis-regulatory elements (Oka et al. 2017; Zhao et al. 2018; Ricci et al. 2019) provided an opportunity to assess the mechanisms and frequency by which TEs may create regulatory variation

In this study, we focused on two potential ways in which TEs might influence the expression of nearby genes; the disruption of regulatory regions and the introduction of novel sequences that may act as regulatory sequences. Insertions into regions of accessible chromatin might be expected to often result in reduced expression of nearby genes or altered patterns of expression. In contrast, TEs that contain accessible chromatin regions may be mobile enhancers that affect expression of both the TE promoter as well as nearby gene promoters. Several studies have found that putative enhancers can be found within transposable elements in the maize genome (Oka et al. 2017; Zhao et al. 2018). We were interested in assessing how frequently the polymorphic insertions could be associated with variable expression for nearby genes to understand the potential for TE polymorphism to generate regulatory diversity. It is worth highlighting the fact that truly assessing the potential for TEs to influence regulation in natural populations may be complicated by the potential fitness consequences of polymorphic TE insertions. If a TE insertion results in significant deleterious or beneficial consequences the allele will likely be a target of selection. Recent studies have found that there are likely many examples of rare deleterious expression states in domesticated maize populations (Kremling et al. 2018) and therefore we monitored both common and rare expression states associated with TE polymorphisms.

### Potential for TEs to reshape chromatin and the epigenome

Active transposition of TEs results in genetic changes including disruption of genes or regulatory elements as well as potential genomic instability due to chromosome breaks or illegitimate recombination. To limit these deleterious events, most genomes have evolved mechanisms to restrict active transposition, including epigenetic silencing through chromatin modifications such as DNA methylation (Hollister and Gaut 2009; Lisch 2013; Springer et al. 2016). This results in highly methylated TEs in plant genomes (Niederhuth et al. 2016) and has been observed to spread outside of the TE sequence to surrounding DNA sequences in some cases (Hollister and Gaut 2009; Eichten et al. 2012; Noshay et al. 2019). As TEs insert into putative regulatory regions, the question becomes not only how the presence of new DNA sequence impacts this region but also the potential for alteration of chromatin patterns. The TE insertion into regions of accessible chromatin can potentially result in loss of accessibility and gains of DNA methylation for the flanking sequences. We observe many examples of TE insertions into accessible chromatin regions for which the regions immediately flanking the TE remain unmethylated and potentially accessible. In some cases, the insertion of a TE within a larger accessible chromatin region results in two smaller accessible chromatin regions on either side of the TE. Often these regions have partial overlap with the edges of the TE. However, there are a subset of examples of TE insertions into accessible regions where the previously accessible and unmethylated regions exhibit high levels of methylation on one or both sides of the TE insertion in the TE-present genotype.

TEs that introduce novel accessible chromatin regions have the challenge of maintaining an unmethylated accessible chromatin region within a highly targeted and condensed repetitive sequence. Even in the TEs that contain an accessible chromatin region, we find that the remainder of the TE is highly methylated. When assessed across three additional genotypes, the methylation state of these accessible chromatin regions was more variable than other unmethylated regions that were outside of TEs. This may suggest that the presence of a TE containing a putative regulatory element in the B73 genome may not predict the presence of an active regulatory element in other genotypes. These would result in the potential for facultative epialleles (Richards 2006; Springer and Schmitz 2017) in which some haplotypes with the TE contain an active regulatory element while others would have a silencer element. This would complicate our ability to make associations between the genetic presence/absence of the TE and the expression level of nearby genes. In our analyses, we made the assumption that when the TE is present the accessible, unmethylated region will be conserved. However, epigenetic polymorphisms would significantly reduce our power. Indeed, careful examination of some examples such as those in figures 6C and D reveal that even though the TE presence is often associated with higher expression for the nearby genes there are some haplotypes that contain the TE but do not show high expression for the nearby gene. These may reflect epigenetic silencing of the regulatory element within these TEs.

### TE influences on regulatory variation for genes

There are massive numbers of polymorphic TE insertions between any two maize genotypes (Wang and Dooner 2006; Springer et al. 2018; Sun et al. 2018; Anderson et al. 2019a). The majority of these polymorphisms likely have little or no impact on gene products or gene expression and are essentially neutral polymorphisms. However, if even a small portion influences gene expression, this could account for a major source of regulatory variation. In this study, we have used chromatin accessibility profiling to narrow the set of TE polymorphisms that might result in altered expression for nearby genes. Specifically, we focused on two classes of polymorphisms that could be assessed based on high quality chromatin accessibility data for the B73 genome (Ricci et al. 2019). The presence of an accessible chromatin region within a TE in B73 enables us to investigate whether the presence of this TE in other maize genotypes is associated with high, or lower, expression of the nearby gene. Alternatively, the presence of an ACR in B73 with a polymorphic TE insertion in PH207, Mo17, or W22 allows for an understanding of how the interruption of an ACR may influence gene expression.

Even in this focused set of TE polymorphisms we find that most of the TE polymorphisms are not significantly associated with altered expression of nearby genes in the tissues we monitored. A majority of genes were found to have little to no change in expression level relative to TE presence/absence (80% of TE-ACRs and 87% of TE insertions into ACRs). This could suggest that these TE-ACRs do not influence expression of the nearby gene. However, it is also possible that in some cases we have not examined the right tissue or growth condition, or that epigenetic instability of the ACR within TEs might complicate our ability to make a genetic association as described above. While the majority of TE polymorphisms were not significantly associated with expression for nearby genes, there are 21 examples of TE insertions into ACRs and 190 examples of TE containing ACRs that are significantly associated with the expression of nearby genes. The lack of strong effects for TE insertions into ACRs was somewhat surprising. In some cases the TE insertions into ACRs may result in dividing a single ACR into two regions separated by the TE. This would predict that there would be instances in the B73 genome in which there are two nearby ACRs that are separated by a TE and the insertion did not necessarily disrupt the functionality of the regulatory region. Interestingly, the examples of TE containing ACRs that are significantly associated with expression are heavily biased towards examples in which the nearby gene is higher expressed. This suggests the TE is providing an enhancer that increases gene expression. In addition to the significant associations, there are also many other examples in which there is substantial variation in expression levels for haplotypes with and without the TE but which lack any statistical significance (outliers). These likely represent examples in which the haplotype with (or without) the TE is rare and only present in one or two genotypes. This might be expected in situations in which TE insertions influence expression resulting in substantial deleterious effects. These outliers are enriched for lower expression of the nearby gene for TE insertions into ACRs but higher expression for the nearby gene for TEs containing ACRs.

A key question we wrestled with in this study, is whether the presence of an ACR within a TE was a property of certain TE families. Given the sequence conservation within TE families, we might predict that the presence of a regulatory element would be conserved in many members of the same TE family. Searching for this consistency is complicated by the focus on uniquely mapping reads. Indeed, we have likely greatly underestimated the number of ACRs within TEs (Figure S4). In many cases, we would only find an ACR in one member of a multi-TE family. These might suggest that the ability to form an accessible region is attributed to both the genetic sequence of the TE as well as local chromatin context. We do find examples of TE families in which there are multiple members with an ACR but even in these families there are other members that lack the ACR (Figure S6-7). In this analysis we do not find strong evidence for TE families in which a common regulatory element is present and accessible for many elements of the same family. This highlights the role for both the DNA sequence of TEs as well as the chromatin landscape of these TEs.

Identification of accessible chromatin regions across the genome has enabled us to narrow in on the ~1% of the genome with potential regulatory function (Rodgers-Melnick et al. 2016; Oka et al. 2017; Zhao et al. 2018; Ricci et al. 2019). By assessing how TE variation could contribute to polymorphisms for these accessible regions we have characterized the potential for TEs to disrupt ACRs or contribute novel ACRs to genes. We assessed both the chromatin and regulatory consequences of these polymorphisms. We find evidence that a subset of TEs containing ACRs are likely providing enhancers to nearby genes. There was little evidence for widespread consequences of insertions of TEs into ACRs. However, many of the TE polymorphisms that strongly influence gene expression might represent rare deleterious alleles. This analysis highlights the potential for TEs to influence gene expression by creating novel expression patterns rather than simply disrupting existing information.

## Acknowledgements

The Minnesota Supercomputing Institute (MSI) at the University of Minnesota provided computational resources that contributed to this research. This work was funded by NSF IOS-1934384 to N.M.S. and C.N.H, NSF IOS-1856627 and NSF IOS-1844427 to R.J.S, and NSF IOS-1543727 to C.N.H. J.M.N. is supported by a Hatch grant from the Minnesota Agricultural Experiment Station (MIN 71-068), R.J.S. is a Pew Scholar in the Biomedical Sciences, supported by The Pew Charitable Trusts. A.P.M is supported by NSF PRFB IOS-1905869.

---

**Figure S1: TE insertions by superfamily.** A) Raw number of TE insertion into ACRs identified (observed) and a control set of random regions of the same size (expected). (B) The proportion of TE insertions into ACRs that are TIRs (purple), LTRs (orange), or Helitrons (green) relative to that expected by chance based on randomized regions of the same size. C) Proportion of DNA transposons that belong to each superfamily for observed (black) or expected based on randomized regions (grey) insertions into ACRs. D) LTR insertions (black) are younger on average than all LTRs in the genome (grey). LTR age is determined by percent identity of the LTR sequences (high % identity represents younger TEs).

**Figure S2: TE insertions split ACRs.** TE insertions into B73 ACRs may result in unmethylated regions on either side of the TE in other genotypes suggesting a TE may split accessible chromatin regions. IGV views display tracks with B73 WGBS methylation (CG blue, CHG green, CHH yellow), B73 ACRs, B73 gene annotations and B73 TE annotations. Each panel identifies a case where a B73 TE is flanked by ACR fragments and the TE is polymorphic in another genotype. A) distal B73 TE absent in PH207, B) proximal B73 TE absent in PH207, C) proximal B73 TE absent in PH207, W22, and Mo17, and D) proximal B73 TE absent in Mo17 and W22,.

**Figure S3: TE-ACR characterization.** A) Proportion of all ACRs in each location category that overlap a TE, majority (>80%) or partial (<80%). Color represents proportion that overlap LTRs (orange), TIRs (purple), or Helitrons (green). B) Distal ACRs are categorized by chromatin pattern as K27me3, Kac, Transcribed, or Unmodified. The proportion of all distal ACRs (grey) and distal ACRs that overlap a TE (black) for each category. C) Proportion of elements containing a distal ACR (>2kb from nearest gene) classified as expressed (evidence for expression across any of the 70 tissues) or silent based on RNA-seq data from Walley et al. Elements were classified by the category of ACR present and the N for each category is shown above each bar.

**Figure S4: ATAC-seq unique and multi-mapping.** A) Proportion of reads uniquely mapped, multi-mapped, or unmapped to the B73v4 genome for an input WGS dataset, ATAC-seq leaf dataset, and ATAC-seq ear dataset. B) Per family unique vs. multi-mapped read counts. Families defined by an ACR (based on unique mapping peak calling) are indicated in red.

**Figure S5: eQTL association.** Posterior probability of association for eQTL with ACRs by chromatin class. Comparison of TE-ACRs (blue) and nonTE-ACRs (purple) to randomized control regions (grey).

**Figure S6: TE-family enrichment for ACRs.** A) Subset of TE families with at least 3 members that have > 30% of their members with an ACR (based on uniquely mapped reads and peak calling). Number above bars indicates TE family size. B) Element age (by percent identity of LTR) for the LTR families with at least 3 members that have > 30% of their members with an ACR

**Figure S7: Sequence similarity across members of the RLX00852 TE family.** VISTA display of sequence similarity for TE family with 3 members containing an ACR (RLX00852Zm00001d00002, RLX00852Zm00001d00003, RLX00852Zm00001d00004) and 2 members lacking an ACR (RLX00852Zm00001d00001 and RLX00852Zm00001d00005). Shown relative to sequence of top TE listed. Grey boxes represent location of ACR in reference sequence.

**Figure S8: Combined dataset TE-Gene expression association.** A/B) Volcano plot of gene expression for genes nearby B73-based ACRs with TE insertions in other genotypes (A) or B73-based TEs containing an ACR (B). A dot is present for each TE-Gene pair for RNA-seq data in each of the 8 tissues. Significant (log2(present/absent) > 2 and q-value < 0.05) and outlier (log2(present/absent) > 5) shown with red and blue points respectively. C/D) Proportion of non-redundant significant (red) or outlier (blue) expression patterns associated with TE-Insertions disrupting an ACR (C) or TE-ACRs (D).

